# Divergent and Convergent TMEM106B Pathology in Murine Models of Neurodegeneration and Human Disease

**DOI:** 10.1101/2024.10.16.618765

**Authors:** Muzi Du, Suleyman C. Akerman, Charlotte M. Fare, Linhao Ruan, Svetlana Vidensky, Lyudmila Mamedova, Joshua Lee, Jeffrey D. Rothstein

**Affiliations:** Department of Neuroscience, Johns Hopkins University School of Medicine, Baltimore, MD, 21205, USA; Brain Science Institute, Johns Hopkins University School of Medicine, Baltimore, MD, 21205, USA; Department of Neurology, Johns Hopkins University School of Medicine, Baltimore, MD, 21205, USA; Department of Psychological and Brain Sciences, Johns Hopkins University Krieger School of Arts and Sciences, Baltimore, MD, 21218, USA

**Keywords:** **Key Terms**: TMEM106B, TAR DNA-binding protein 43, neurodegenerative disease, amyotrophic lateral sclerosis, frontotemporal dementia, Alzheimer’s disease, tauopathy, murine models of neurodegenerative disease, pathology

## Abstract

TMEM106B is a lysosomal/late endosome protein that is a potent genetic modifier of multiple neurodegenerative diseases as well as general aging. Recently, TMEM106B was shown to form insoluble aggregates in postmortem human brain tissue, drawing attention to TMEM106B pathology and the potential role of TMEM106B aggregation in disease. In the context of neurodegenerative diseases, TMEM106B has been studied *in vivo* using animal models of neurodegeneration, but these studies rely on overexpression or knockdown approaches. To date, endogenous TMEM106B pathology and its relationship to known canonical pathology in animal models has not been reported. Here, we analyze histological patterns of TMEM106B in murine models of *C9ORF72*-related amyotrophic lateral sclerosis and frontotemporal dementia (C9-ALS/FTD), SOD1-related ALS, and tauopathy and compare these to postmortem human tissue from patients with C9-ALS/FTD, Alzheimer’s disease (AD), and AD with limbic-predominant age-related TDP-43 encephalopathy (AD/LATE). We show that there are significant differences between TMEM106B pathology in mouse models and human patient tissue. Importantly, we also identified convergent evidence from both murine models and human patients that links TMEM106B pathology to TDP-43 nuclear clearance specifically in C9-ALS. Similarly, we find a relationship at the cellular level between TMEM106B pathology and phosphorylated Tau burden in Alzheimer’s disease. By characterizing endogenous TMEM106B pathology in both mice and human postmortem tissue, our work reveals considerations that must be taken into account when analyzing data from *in vivo* mouse studies and elucidates new insights supporting the involvement of TMEM106B in the pathogenesis and progression of multiple neurodegenerative diseases.

## Introduction

Neurodegenerative diseases are a notoriously enigmatic class of disorders, with varied clinical and cellular presentation. One common pathological feature of neurodegenerative disorders, however, is the misfolding and aggregation of proteins such as TAR DNA-binding protein 43 (TDP-43), amyloid-β, tau, or α-synuclein [31, 70]. These misfolded proteins are thought to be toxic to cells, and have been shown to elicit neurodegeneration *in vitro* and in animal models [70]. As such, protein aggregates are frequently studied as potential biomarkers of disease [44] or as therapeutic targets [83]. Unfortunately, although protein aggregation is observed across neurodegenerative diseases, the identity of the specific aggregated protein(s) varies, complicating efforts toward pharmacological intervention [70]. Recently, <u>T</u>rans<u>mem</u>brane protein 106B (TMEM106B) aggregates were described in the postmortem brain tissues of patients with a wide range of neurodegenerative diseases, including Alzheimer’s Disease (AD) [75], Parkinson’s Disease (PD) [19, 75], frontotemporal lobar degeneration (FTLD) [38, 75], amyotrophic lateral sclerosis (ALS) [9, 75], as well as other neurodegenerative diseases and normal aging [9, 75].

TMEM106B was originally identified as the most significant risk factor for FTLD with TDP-43 inclusions (FTLD-TDP) [60, 90]. Since then, several single nucleotide polymorphisms (SNPs) of TMEM106B have been identified as modifiers of disease phenotypes in frontotemporal dementia (FTD) [26, 30, 87]. In one case study, homozygosity of the TMEM106B protective allele (rs3173615) completely shielded autosomal dominant progranulin (GRN) mutation carriers from developing FTD [63]. This result suggests that TMEM106B is involved in affecting disease penetrance.

In addition to being a disease modifier for FTD, several genome-wide association studies (GWAS) have uncovered TMEM106B variants that are relevant to other neurodegenerative diseases. For instance, one risk variant (rs1990622) is implicated in the pathologic presentation of AD [10, 47, 72, 90], and genetic editing of the risk allele to a protective allele of TMEM106B rescued cognitive decline and neurodegeneration in an animal model of tauopathy [18]. Furthermore, TMEM106B variants are significantly correlated with the transcriptional signature of biological aging [69], cognition [92], brain volume [1], and levels of neuronal markers and neuronal proportion [67] specifically in aged cohorts without clinical diagnosis of dementia.

TMEM106B is a type 2 lysosomal/late endosomal membrane protein that is highly expressed in the brain, particularly in neurons and oligodendrocytes [21, 22, 97]. Several studies have found that TMEM106B plays an important role in regulating lysosome size [5, 10, 80], axonal transport of lysosome [50, 76, 80], lysosomal acidification [10, 22, 45, 97], lysosomal protein homeostasis [22, 42, 50], and autophagy [22, 50]. In agreement with these observations, a recent study found that loss of TMEM106B in an animal model of tauopathy results in increased cytoskeletal disruption, impaired autophagy, errors in lysosomal trafficking along the axon, and enhanced gliosis [20].

To date, studies investigating TMEM106B in animal models have introduced genetic modulation of the *TMEM106B* gene to generate knockdown/out or overexpression models, all of which could produce non-physiological artifacts. However, studying TMEM106B at endogenous levels in existing mouse models for neurodegeneration has not yet been performed. Mouse models are a valuable resource in the field of neurodegeneration, as *in vivo* studies allow for relatively facile temporal and cellular genetic manipulation in complex organisms. Moreover, the murine proteome shares high sequence similarity with humans, allowing for novel insights into human biology [15]. Thus, understanding how mouse models relate to human pathology may be important for interpreting mouse studies. Here we characterize TMEM106B phenotypes in three different models of neurodegeneration and compare these results to disease-matched human postmortem tissue. Indeed, we find that there are important differences between what is observed in human tissue and mouse tissue. Importantly, we also find convergent phenotypes that reveal potentially novel biological contributions of TMEM106B to TDP-43 nuclear clearance in C9-ALS and tau pathology in Alzheimer’s disease. Taken together, these results further suggest an important role of TMEM106B in neurodegenerative diseases and provide new insights into the cellular processes associated with the aberrant TMEM106B pathology in both murine models and human diseases.

## Results

### TMEM106B Forms Cytoplasmic Inclusions in an AAV-based Mouse Model of C9-ALS

The GGGGCC (G_4_C_2_) hexanucleotide expansion in intron 1 of the *C9ORF72* gene is the most common genetic cause of ALS (C9-ALS) [68]. The presence of the G_4_C_2_ intron expansion is thought to lead to three, non-mutually exclusive, pathological events: (1) haploinsufficiency of the C9orf72 protein, (2), the expression and accumulation of toxic repeat RNA species, and (3), the accumulation of toxic dipeptide repeat (DPR) proteins produced via repeat-associated non-AUG (RAN) translation [56, 68]. Many groups have established mouse models to study C9-ALS *in vivo*, including both adeno-associated virus (AAV)-based and bacterial artificial chromosome (BAC)-based expression. The number of disease-associated repeats being expressed in these models ranges from 66 [13] to over 400 [37, 48], as compared to humans, in which the pathological number of repeats identified in C9-ALS patients ranges from ∼30 to over 4,000 [27].

Previous studies have identified TMEM106B as a genetic modifier of disease penetrance of *C9ORF72* expansion carriers [26, 89]. To investigate endogenous TMEM106B in an *in vivo* model of C9-ALS, we employed an AAV-based approach utilizing an AAV vector harboring either 2 or 149 G_4_C_2_ repeats [11]. For these experiments, mice are given an intracerebroventricular injection of AAV vector containing (G_4_C_2_)_149_, or (G_4_C_2_)_2_ at P0. Previous reports with these animals found that by 6 months, the expression of (G_4_C_2_)_149_ leads to behavioral defects, neuronal loss, increased gliotic GFAP reactivity, and the accumulation of RNA foci, phosphorylated TDP-43 inclusions, and DPR aggregates in the motor cortex [11]. At 12 months, animals expressing (G_4_C_2_)_149_, but not (G_4_C_2_)_2_, also show inclusions of several stress granule-related proteins, which colocalize with DPRs and phosphorylated TDP-43 [11]. Thus, this model recapitulates many of the hallmark features of C9-ALS pathology.

For 9-month-old animals that had been injected with either (G_4_C_2_)_149_, or (G_4_C_2_)_2_, we first stained mouse brain sections using a commercially available antibody (TMEM-Sigma) that was previously shown to detect TMEM106B aggregates in human postmortem brain tissue [53, 64], and recognizes sequences homologous with the murine TMEM106B gene. By both DAB (**Figure 1A, B**) and immunofluorescence staining (**Figure 1C-E**) we observed a distinct pattern of TMEM106B perinuclear inclusions that were specifically enriched in animals injected with AAV-(G_4_C_2_)_149_, and not in the control AAV-(G_4_C_2_)_2_ animals.

**Figure 1:**
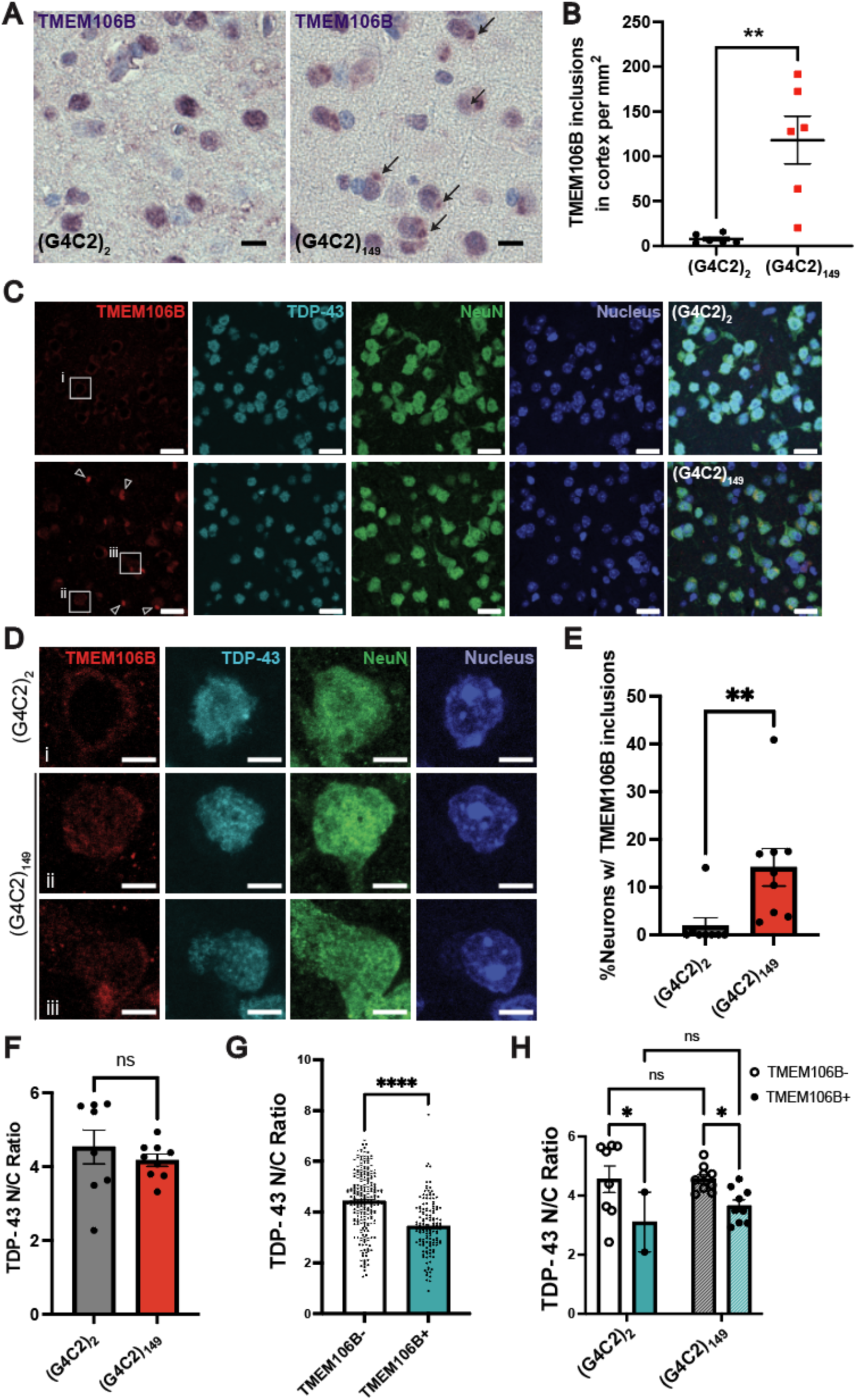
TMEM106B forms neuronal perinuclear inclusions that are associated with decreased nuclear TDP-43 in (G_4_C_2_)_149_ repeat expressing mice. (A) Representative images of cortex with DAB staining against TMEM106B in mice injected with either (G_4_C_2_)_2_ or (G_4_C_2_)_149._ Arrows indicate observed TMEM106B positive perinuclear inclusions. Scale bar = 10 µm. (B) Quantification of TMEM106B inclusions in the cortex ((G_4_C_2_)_2_ n = 6 and (G_4_C_2_)_149_ n = 6). Dots represent individual animals, bars represent means ± SEM. Student’s unpaired t-test, p = 0.001. (C) Immunofluorescence co-staining of TMEM106B, TDP-43 and NeuN (a neuronal marker) in the motor cortex from AAV-(G_4_C_2_)_2_ (upper panel) or AAV-(G_4_C_2_)_149_ (lower panel) injected mice. Arrowheads indicate TMEM106B perinuclear inclusions that are enriched in (G_4_C_2_)_149_ mice. Enlarged images of representative cells are outlined and shown in panel D. Scale bar = 20 µm. (D) Zoomed in images of individual cells showing an example neuron with a TMEM106B perinuclear inclusion (cell iii) which has a disrupted TDP-43 nuclear-to-cytoplasmic (N/C) ratio. Scale bar = 5 µm. (E) Quantification of percentage of neurons with TMEM106B perinuclear inclusions in (G_4_C_2_)_2_ (n = 8) and (G_4_C_2_)_149_ (n = 9) mice. Mann-Whitney test, p = 0.0012. (F) Quantification of averaged TDP-43 N/C ratio in (G_4_C_2_)_2_ (n = 8) and (G_4_C_2_)_149_ (n = 9) mice. Dots represent individual animals, bars represent means ± SEM. Students t-test was used to compare groups. (G) Quantification of TDP-43 N/C ratio in neurons with TMEM106B perinuclear inclusion (TMEM106B+, n = 140) and neighboring neurons in the same tissues that do not have an inclusion (TMEM106B-, n = 232) across all (G_4_C_2_)_2_ (n = 8) and (G_4_C_2_)_149_ (n = 9) mice. Mann-Whitney test, p<0.0001. Dots represent cells, bars represent means ± SEM. (H) Quantifying the TDP-43 N/C ratio for either (G_4_C_2_)_2_ or (G_4_C_2_)_149_ mice and comparing the average TDP-43 N/C ratio per animal by TMEM106B phenotype shows that there is a decreased N/C ratio in cells with TMEM106B cytoplasmic inclusions. Dots represent averages for each phenotype for each animal. Bars represent means ± SEM. Two-way ANOVA with multiple comparisons; p = 0.0419 for TMEM106B-vs TMEM106B+ in the (G_4_C_2_)_2_ group; p = 0.0329 for TMEM106B-vs TMEM106B+ in the (G_4_C_2_)_149_ group; ns, p>0.05.

Next, we stained the AAV-injected mouse tissues with a separate published antibody (antibody TMEM239) that has been shown to recognize TMEM106B C-terminal aggregates [3, 75] and whose epitope has exact sequence homology with the murine TMEM106B gene. Unlike the large perinuclear inclusions seen using the TMEM-Sigma antibody, TMEM239 immunoreactivity revealed a morphology which was more punctate (**Figure S1A**). Notably, animals injected with AAV-(G_4_C_2_)_149_ developed a significantly greater number of intracellular TMEM239-positive puncta compared to the (G_4_C_2_)_2_ control animals (**Figure S1B, C**).

To further characterize the TMEM106B inclusions observed by TMEM-Sigma antibody staining, we next co-stained TMEM106B with markers for autophagy (p62), stress granules (eukaryotic initiation factor 3η, eIF3η), and lysosomes (cathepsin D, CthD). Interestingly, TMEM106B does not colocalize strongly with any of these markers (**Figure S1D, E**), suggesting that these TMEM-Sigma-positive structures do not reflect the canonical function of TMEM106B and may be linked to other pathological changes related to (G_4_C_2_)_149_ expression.

Genetic variants of TMEM106B[53, 90] and the deposition of TMEM106B C-terminal fragments[53] were each previously shown to be associated with pathological TDP-43 inclusions. However, given that TMEM106B aggregation occurs in multiple neurodegenerative diseases as well as in general aging[9, 75], whether TMEM106B C-terminal aggregates correlate with TDP-43 pathology specifically in ALS is unclear. Thus, to investigate the relationship between the TMEM106B inclusions observed in C9 animals and changes in TDP-43 at the cellular level, we co-stained tissues for TMEM106B and TDP-43 (**Figure 1C, D**).

Studies have shown that TDP-43 pathology in ALS is defined by its nuclear clearance, rather than TDP-43 aggregation which only occurs in a small portion of neurons [17, 71, 94]. Therefore, we measured the TDP-43 nuclear-to-cytoplasmic (N/C) ratio in the motor cortices of AAV-injected mice. On average, the TDP-43 N/C ratio is not significantly different in AAV-(G_4_C_2_)_149_ mice compared to AAV-(G_4_C_2_)_2_ mice (**Figure 1F**). However, we found that the specific sub-group of neurons with TMEM106B perinuclear inclusions has a significantly lower TDP-43 N/C ratio compared to neighboring neurons without TMEM106B inclusions (**Figure 1G, H**) suggesting a relationship between abnormal TMEM106B inclusion formation and altered TDP-43 cellular distribution. Overall, these results reveal a previously unreported TMEM106B pathology characterized by a perinuclear inclusion. In addition, the enrichment of TMEM106B inclusions in (G_4_C_2_)_149_ animals and its correlation with TDP-43 nuclear clearance *in vivo* could suggest that TMEM106B may relate, in as yet an undefined mechanism, to disease pathogenesis in C9-ALS.

### Cytoplasmic TMEM106B Puncta Coincide with Reduced Nuclear TDP-43 in Human C9-ALS and C9-ALS/FTD Tissue

To investigate whether our observations in mouse tissue were representative of human disease, we next performed immunohistochemistry on human motor and occipital cortex from either healthy control or C9-ALS and C9-ALS/FTD patients (**Table 3**). Using the TMEM-Sigma antibody, we did not detect perinuclear inclusions in human tissue (**Figure 2A, B, S2A, B**). We also did not observe global differences in the TDP-43 N/C ratio by disease status (**Figure 2C, S2C**). However, we did observe intracellular puncta in the motor cortex that resemble what has been reported before in FTLD-TDP tissues (**Figure 2A, D**) [53, 64].

**Figure 2:**
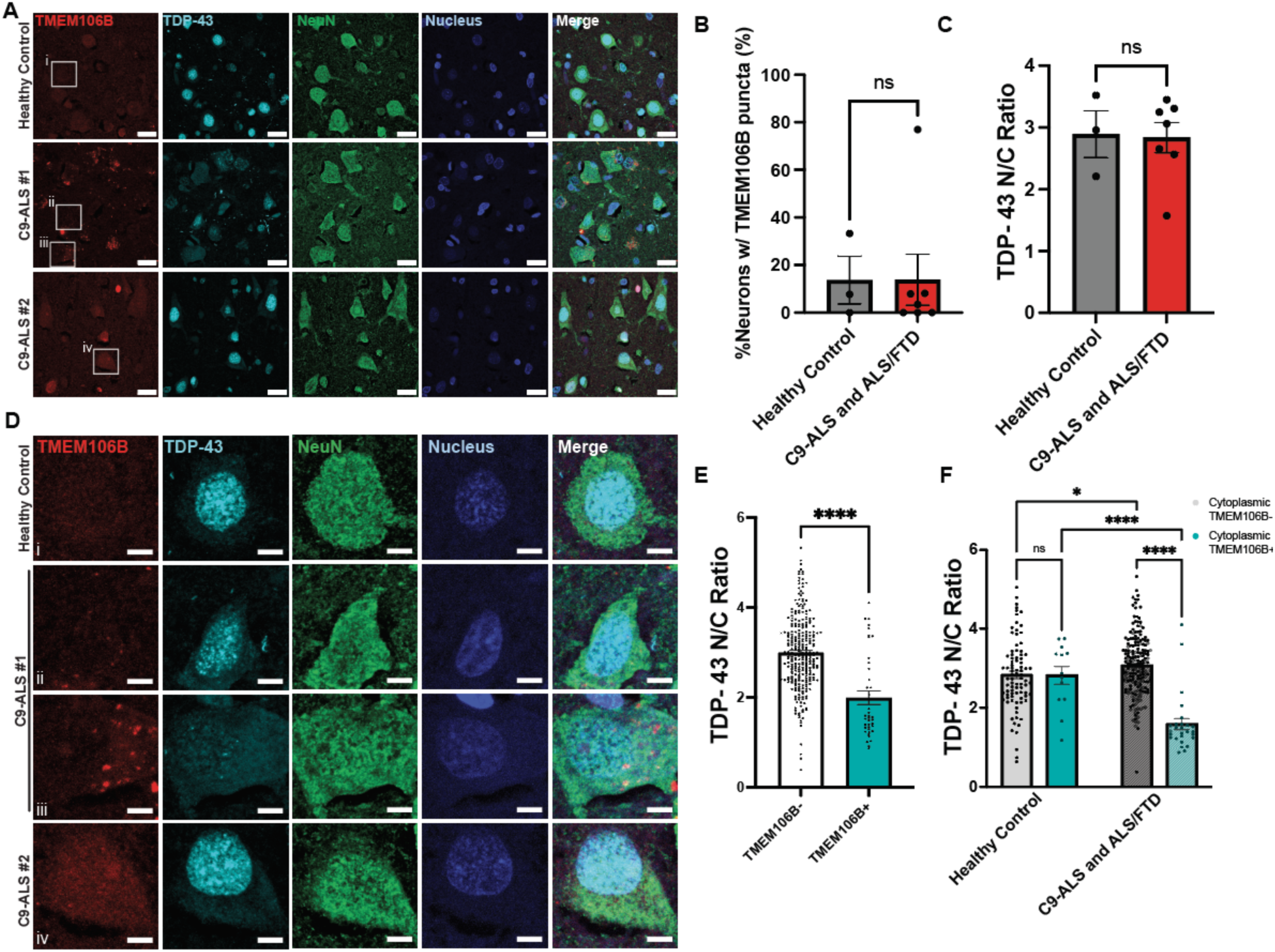
Cytoplasmic TMEM106B punctate is associated with decreased nuclear TDP-43 in the motor cortices of human C9-ALS and ALS/FTD cases. **(A)** Human motor cortex co-stained with TMEM-Sigma antibody, TDP-43 and NeuN. One neurologically healthy control, one C9-ALS patient (#1) with severe TDP-43 nuclear clearance, and one C9-ALS patient (#2) with relatively intact TDP-43 localization are shown. Enlarged images of representative cells are outlined and shown in Figure (D). Scale bar = 20 µm. **(B)** Quantification of the percentage of neurons with intracellular TMEM106B puncta from healthy control (n = 3), and C9-ALS and ALS/FTD (n = 7) patients. Dots represent individual people, bars represent means ± SEM. Mann-Whitney test, p = 0.9333. **(C)** Quantification of averaged TDP-43 nuclear to cytoplasmic (N/C) ratio from healthy control (n = 3), and C9-ALS and ALS/FTD (n = 7) patients. Dots represent individual people, bars represent means ± SEM. Unpaired t-test, p = 0.9. **(D)** Zoomed in images of individual cells showing an example neuron with TMEM106B cytoplasmic puncta (cell iii) with severe TDP-43 nuclear clearance. Scale bar = 5 µm. **(E)** Quantification of TDP-43 N/C ratio in neurons with TMEM106B cytoplasmic puncta (TMEM106B+, n = 40) and those without (TMEM106B-, n = 280) across healthy control (n = 3) and C9-ALS and ALS/FTD (n = 7) patients. Dots represent individual cells, bars represent means ± SEM. Mann-Whitney test, p<0.0001. **(F)** Quantification of the TDP-43 N/C ratio of individual neurons with or without TMEM106B cytoplasmic puncta from healthy controls or C9-ALS and ALS/FTD patients grouped by both patient diagnosis and TMEM106B phenotype. For the C9-ALS and ALS/FTD bars, closed dots represent C9-ALS, and open diamonds represent C9-ALS/FTD. Data points represent individual cells, bars represent means ± SEM. A two-way ANOVA with multiple comparisons test was performed; *, p = 0.024; ****, p<0.0001.

To investigate whether there was a relationship between the presence of TMEM106B puncta and TDP-43 distribution as we observed in mice, we next quantified the N/C ratio of TDP-43 based on TMEM106B phenotype (**Figure 2E, F**). Indeed, in human motor cortex, neurons that contain cytoplasmic TMEM106B puncta displayed a significantly reduced TDP-43 N/C ratio (**Figure 2E**). Interestingly, subcategorization of neurons into healthy and diseased groups reveals that the overall reduction of nuclear TDP-43 in the TMEM106B puncta-positive cells is specific to the disease group (**Figure 2F**). That is, patients with C9-ALS/FTD show a TMEM106B-related decrease in nuclear TDP-43. In addition, we found that the presence of neuronal TMEM106B puncta is rare in the occipital cortices for both healthy and C9 patients (**Figure S2A, B**), suggesting that the TMEM106B:TDP-43 correlation is specific to the affected brain region in C9-ALS and C9-ALS/FTD. In line with previous studies on the association between TMEM106B genetic variants and TDP-43 aggregation pathology [53, 90], our data provide new evidence that TMEM106B could be related to the nuclear clearance of TDP-43 specifically in C9-ALS and C9-ALS/FTD at the cellular level.

### TMEM106B Does Not Form Inclusions in a SOD1 ALS Model

We next sought to characterize another genetic model of ALS to see if our findings in the C9-ALS mouse model generalize to other forms of ALS. Thus, we chose the SOD1 G93A mouse model, which has long been used to study ALS *in vivo* [29, 88]. In this model, mice transgenically express mutant hSOD1^G93A^, resulting in neurofilament aggregation, loss of motor neurons, and astrocytosis by 3 months of age, followed by progressive paralysis and premature death [29, 88]. Although no studies have yet described TMEM106B aggregation in SOD1-ALS specifically, pathological misfolded SOD1 impacts autophagic processes [40, 57, 84, 96], which could affect or be affected by TMEM106B aggregation. Additionally, TDP-43 cytoplasmic inclusions are largely absent in SOD1-ALS patients [51, 61, 86], indicating a pathologically distinct mechanism of neurodegeneration.

First, we established SOD1 pathology in this model by staining for SOD1 in the motor cortex, hippocampus, midbrain, and hindbrain of 3-month-old animals (**Figure 3A, S3A**). As expected, we saw that SOD1 is highly expressed in transgenic animals, and not in control animals. Moreover, C4F6, a well-characterized antibody for misfolded SOD1 that detects an exon 4 epitope (**Table 1**), shows positive staining in the motor cortex and hippocampus of transgenic animals (**Figure S3A**), in line with reports in both human and mouse tissue [23, 41, 62, 65, 66]. We also saw Iba1+ staining in the midbrain and hindbrain, indicative of microgliosis (**Figure 3B**). Although SOD1 staining was present throughout the brain, we mainly observed vacuolization in the midbrain and hindbrain (**Figure 3A, C**), consistent with previous reports of vacuolar degeneration in models of SOD1 ALS [28, 33, 73, 93]. Indeed, by both DAB staining (**Figure 3A**) and immunofluorescence staining (**Figure 3C**), we observed robust vacuolization in the midbrain and hindbrain specifically in transgenic animals. Thus, these animals display the pathological features expected of the SOD1 ALS mouse model.

**Figure 3:**
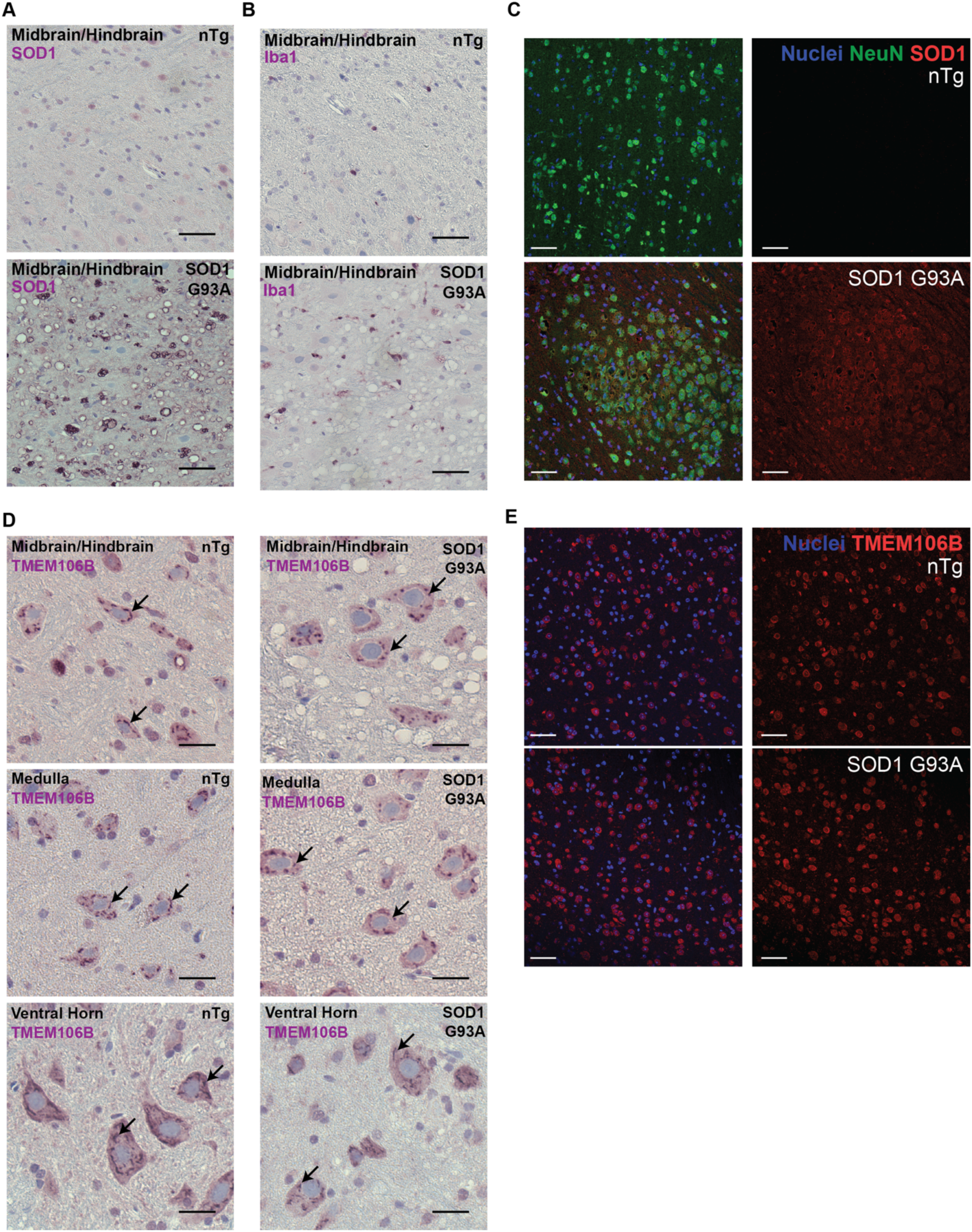
TMEM106B does not have altered distribution in mice expressing ALS-mutant G93A SOD1. **(A)** Midbrain/hindbrain region in non-transgenic (nTg) animals and animals expressing G93A SOD1 with DAB staining against SOD1. Vacuolization is apparent in transgenic animals. Scale bar = 25 µm. **(B)** Representative images of midbrain/hindbrain region in nTg and G93A SOD1 animals with DAB staining against Iba1 showing increased Iba1 reactivity in transgenic animals, as well as vacuolization. Scale bar = 25 µm. **(C)** Performing immunofluorescence confocal microscopy on the midbrain of either non-transgenic or SOD1 G93A transgenic animals shows clear overexpression of SOD1 in the transgenic animals and vacuolization. Images are representative of n = 8 nTg and 7 transgenic animals. Scale bar = 50 µm. **(D)** Representative images of midbrain/hindbrain, medulla and ventral horn region with DAB staining against TMEM106B. Scale bar = 25 µm. A punctate like staining pattern as shown by arrows in both nTg and transgenic animals. **(E)** Immunofluorescence confocal microscopy for TMEM106B does not reveal any overt differences in TMEM106B staining between non-transgenic and transgenic animals. Images are representative of n = 8 nTg and 7 transgenic animals. Scale bar = 50 µm.

**Table 1.** List of antibodies for tissue staining.

| Target | Source | Catalog # | Dilution |
| --- | --- | --- | --- |
| TMEM106B | Sigma | SSAB2106773 | IF: 1:500 |
| TMEM106B (TMEM239) | Gift from Michel Goedert |  | IF: 1:500 |
| NeuN | Millipore | ABN91 | IF: 1:500 (mouse);<br>1:100 (human) |
| SOD1 | Abcam | Ab52950 | IF: 1:50 |
| Misfolded SOD1 | MédiMabs | MM-0070-2-P | IF: 1:50 |
| AT8 | ThermoFisher | MN1020 | DAB: 1:250<br>IF: 1:500 |
| p62 | BD Biosciences | 610832 | IF: 1:200 |
| eIF3η | Santa Cruz | sc-137214 | IF: 1:100 |
| Cathepsin D | B&D Systems | MAB1029 | IF: 1:100 |
| TDP-43 | Abcam | ab104223 | IF: 1:500 |
| Goat anti-Chicken IgY (H+L) Secondary Antibody, Alexa Fluor 488, Invitrogen | Invitrogen | A-11039 | IF: 1:1000 |
| Goat Anti-Mouse IgG Polyclonal Antibody (CF™ 647) | Biotium | 20281-1 | IF: 1:1000 |
| Goat anti-Rabbit IgG (H+L) Highly Cross-Adsorbed Secondary Antibody, Alexa Fluor™ 568 | ThermoFisher | A-11036 | IF: 1:1000 |

**Table 2.** List of key reagents and resources.

| Reagent or Resource | Source | Catalog # |
| --- | --- | --- |
| Hoechst 33342 | BD Biosciences via GRCF | 561908 |
| DAPI | Invitrogen | D1306 |
| ProLong Gold Antifade Mountant | Thermo Fisher | P36930 |
| 18x18 mm High Tolerance Coverslips | MatTek | PCS-170-1818 |
| DAKO Envision+HRP polymer kits | Agilent | K4003 and K4001 |
| DAKO Serum-Free Protein Block | Agilent | X0909 |
| DAKO antibody Diluent | Agilent | S3022 |
| ImmPACT® VIP Substrate Kit | Vector Laboratories | SK-4605 |
| Hematoxylin QS Counterstain | Vector Laboratories | H-3404-100 |
| Aqua-Poly/Mount | Polysciences | 18606-20 |
| ZEN Microscopy Software | Zeiss | Zen Ver. 3.5 |
| CellProfiler | [81] | N/A |
| Prism 10 | GraphPad | N/A |
| ImageJ (FIJI) | [74] | N/A |

After demonstrating SOD1-relevant pathology, we next examined TMEM106B localization in the midbrain and hindbrain, medulla, and ventral horn. By DAB staining, we observed punctate-like TMEM106B staining in both non-transgenic and transgenic mice (**Figure 3D**). Similarly, using immunofluorescence, we did not observe any notable difference between TMEM106B staining in the midbrain of non-transgenic and transgenic mice (**Figure 3E**). Moreover, we did not observe the large cytoplasmic inclusions found in the AAV-C9-ALS model using either DAB staining (**Figure 3D**) or immunofluorescence staining (**Figure 3E**).

Because one key pathological feature of the SOD1 mice is vacuolization [28, 33, 73, 93], and because TMEM106B is a membrane-bound protein, we wondered whether TMEM106B localizes to the vacuolar structures formed in the midbrain and hindbrain. Thus, we examined high-resolution images of cells with large vacuoles. However, we did not observe an increase in TMEM106B staining around the vacuole perimeter (**Figure S3B**). We also did not observe any changes to TMEM106B localization in the motor cortex, despite the positive staining for misfolded SOD1 (**Figure S3A, C**). Overall, these results indicate TMEM106B pathology is not a prevalent histological feature of SOD1-ALS and suggest that TMEM106B may play a role in the pathogenesis of specific forms of ALS, such as those caused by C9orf72 repeat expansion.

### TMEM106B Immunoreactivity Correlates with Accumulation of Phosphorylated Tau at the Early Stages of the PS19 Mouse Model of Tauopathy

TMEM106B is not only a genetic risk factor for ALS and FTD [52, 90] but is also implicated in other neurodegenerative diseases such as Alzheimer’s disease [24, 34, 35]. Additionally, genetic manipulation of TMEM106B expression in murine models of tauopathy have revealed a potentially significant role of TMEM106B in tau-related neurodegenerative diseases [18, 20]. However, a rigorous analysis of the histological pattern of endogenous TMEM106B in early- and late-stage murine models of tauopathy has not yet been conducted. Thus, we next looked at the PS19 mouse model, a well-characterized *in vivo* system for studying tauopathy [95].

PS19 mice transgenically express the FTD with parkinsonism linked to chromosome 17 (FTDP-17)-associated P301S mutant of tau[95]. By 3 months of age, PS19 mice begin to display a motor phenotype, leading to paralysis by 7-10 months, with a median survival of 9 months [95]. Further characterization of these animals has established that PS19 mice accumulate insoluble tau and phosphorylated tau (pTau) [95]. This pathology is accompanied by neuronal loss in the hippocampus and brain atrophy [95]. Only ∼20% of PS19 animals survive to 12 months of age, at which point there is significant loss of brain volume [95]. Thus, 12-month-old PS19 animals represent late-stage tauopathy.

Consistent with the established phenotype of the PS19 model, we observe robust accumulation of pTau in the hippocampus and motor cortex of 12-month-old PS19 animals (**Figure S4A**) by DAB staining with the Ser202/Thr205 phosphorylation-dependent tau antibody, AT8 (**Table 1**). Using immunofluorescence to quantify pTau in hippocampal neurons of the dentate gyrus, we find both significant loss of NeuN+ neurons and a significant increase in pTau in PS19 animals relative to non-transgenic controls (**Figure S4B**).

Next, we performed DAB staining of TMEM106B in the hippocampus and did not find any obvious differences in TMEM106B staining between non-transgenic mice and PS19 animals (**Figure S4C**). Similarly, we did not observe a significant difference in immunofluorescence reactivity for TMEM106B, although there was a slight increase in TMEM106B signal for PS19 animals (**Figure S4D, E**). This is intriguing, as previous reports indicate that the TMEM106B rs1990622-A variant that is associated with increased risk for Alzheimer’s disease is also correlated with higher levels of the TMEM106B protein in the hippocampus [25, 47, 59].

Although clearly distinct from the inclusions formed in the AAV-C9 tissue, we did observe some areas of high TMEM106B reactivity in the PS19 tissue. When we co-stained these tissues with TMEM106B and AT8, we found that there was no significant correlation between AT8 and TMEM106B staining intensities in the 12-month-old animals (**Figure S4F**). Indeed, closer examination of hippocampal neurons with pTau inclusions and TMEM106B reactivity showed that there was no co-localization between the two structures, consistent with previous work showing that TMEM106B-positive species do not co-localize with Tau [64] (**Figure S4G**).

We next wondered whether there was any distinct pathological presentation in the PS19 model at 9 months of age, when neuronal loss is not as severe [95]. Indeed, immunofluorescence staining of NeuN in the hippocampus at 9 months of age reveals that the loss of NeuN+ cells in the dentate gyrus is only modestly significant in PS19 animals relative to non-transgenic controls (**Figure 4A, B**). However, 9-month-old PS19 animals are still robust models of tauopathy, as pTau staining is significantly elevated compared to control (**Figure 4C**). Surprisingly, whereas there was a slight but not significant increase in TMEM106B immunoreactivity at 12 months for the PS19 cohort, in the 9-month-old animals this increase is significant (**Figure 4D**). Moreover, there is a significant positive correlation between the intensity of AT8 staining and TMEM106B immunoreactivity (**Figure 4E**). Thus, at earlier time points, when neurons have not yet died, TMEM106B and pTau both show increased immunoreactivity, which may reflect underlying pathological changes.

**Figure 4:**
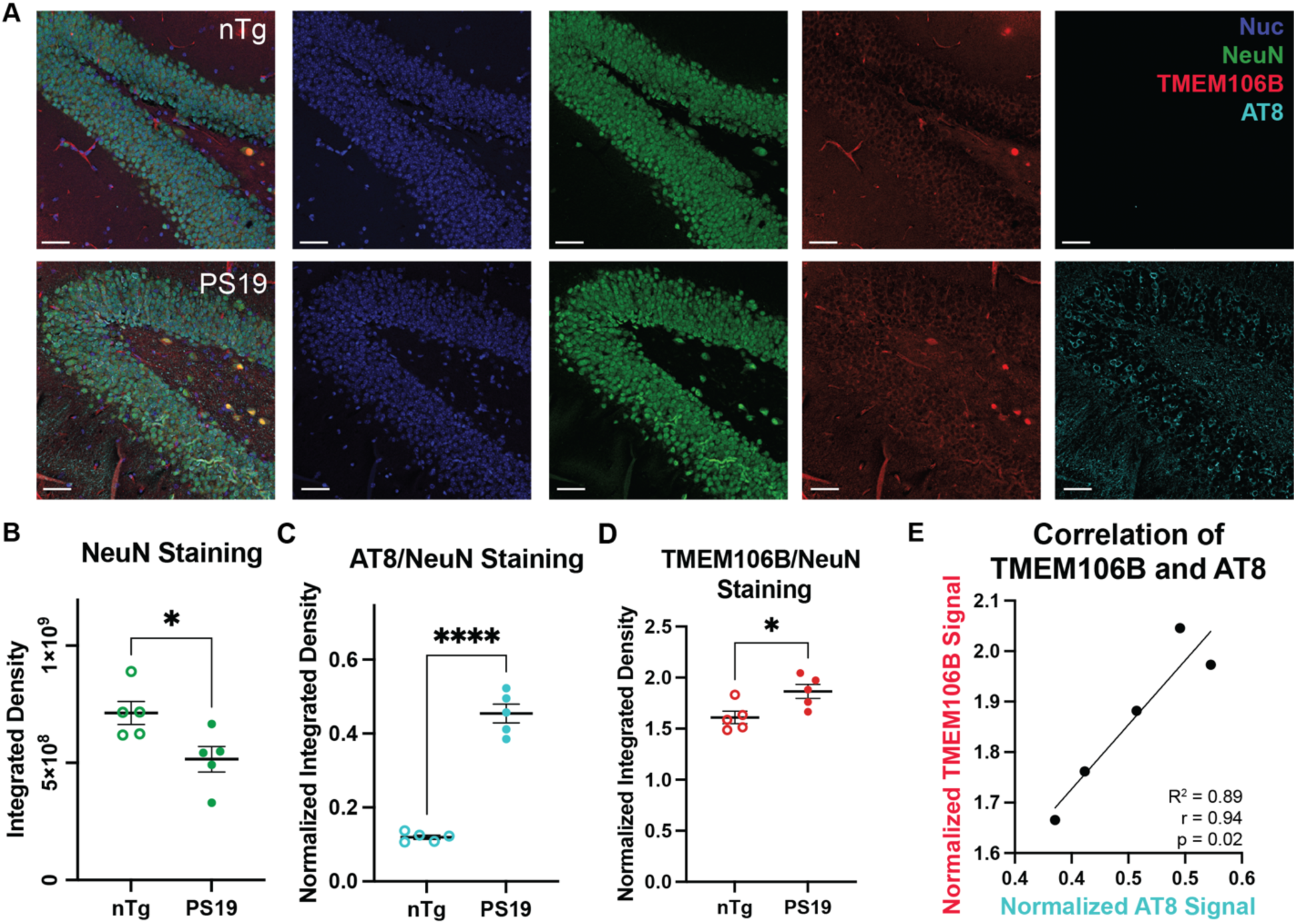
Correlation of TMEM106B and phosphorylated tau is observed in 9-month-old PS19 mice. **(A)** Representative immunofluorescence images of the hippocampal dentate gyrus region of non-transgenic (nTg) and PS19 mice stained for NeuN, TMEM106B, and phosphorylated Tau (AT8). Quantification of NeuN **(B)**, TMEM106B **(C)**, and AT8 **(D)** staining in 9-month-old mice show that there is a significant difference in the signal intensity between transgenic and nTg mice. The signal for TMEM106B and AT8 was normalized to the signal of its corresponding NeuN channel. Each data point represents an individual image from n = 5 each of nTg and PS19 animals. Bars represent means ± SEM. A student’s t-test was used to compare nTg and transgenic animals; *, p<0.03; ****, p<0.0001. **(E)** Linear regression and correlation analysis between TMEM106B and AT8 signals.

### TMEM106B Punctate-Like Structures in AD Human Tissue Correlate but not Colocalize with Phosphorylated Tau

Our results in the PS19 model suggest that there may be co-pathology between TMEM106B and the accumulation of phosphorylated tau. Thus, we next investigated TMEM106B phenotypes in human tauopathy using two different disease cohorts: AD and AD with limbic-predominant age-related TDP-43 encephalopathy (AD/LATE). AD/LATE can be associated with a higher tau burden [85], thereby providing additional insight into the relationship between tau and TMEM106B in disease.

We first analyzed TMEM106B deposition in postmortem tissue from control and AD patients using DAB (**Table 4**). We focused on the cornu ammonis (CA) region of the hippocampus, as this area is known to be heavily affected in AD [4]. In control samples, TMEM106B is diffused throughout the cell, with both cytoplasmic and nuclear staining by DAB (**Figure S5A**). In AD tissue, by contrast, TMEM106B forms aggregated puncta (**Figure S5B**). This agrees with previous studies which have shown that TMEM106B forms neuronal aggregates patients with AD and other tauopathies [64].

Next, we performed co-staining of TMEM106B and pTau in postmortem tissue from patients with AD and AD/LATE (**Figure 5A**, **Table 4**). As compared to control samples, histologically defined AD and AD/LATE patient tissues had significantly higher levels of NeuN-normalized AT8 staining in the hippocampus, indicative of the accumulation of pTau (**Figure 5B**). Quantification of TMEM106B immunoreactivity showed that the levels of TMEM106B are not increased in AD tissue but are increased in AD/LATE patients (**Figure 5C**). Previous reports have shown that the risk allele rs1990622 is associated with higher levels of TMEM106B mRNA and protein [10, 47, 90], however as the genomic information of the patients is not available to us, it is unknown whether the patients characterized here are carriers of this risk variant. Interestingly, we identified a subpopulation of cells in AD/LATE patients that had higher TMEM106B staining (**Figure 5C**). Thus, to determine whether TMEM106B levels were related to pTau burden in human disease, we calculated the correlation between TMEM106B and AT8 intensities for control and disease cohorts (**Figure 5D**). We find that there is a slight but significant correlation between TMEM106B and pTau levels in AD and AD/LATE tissues, with AD/LATE patients showing the strongest correlation.

**Figure 5.**
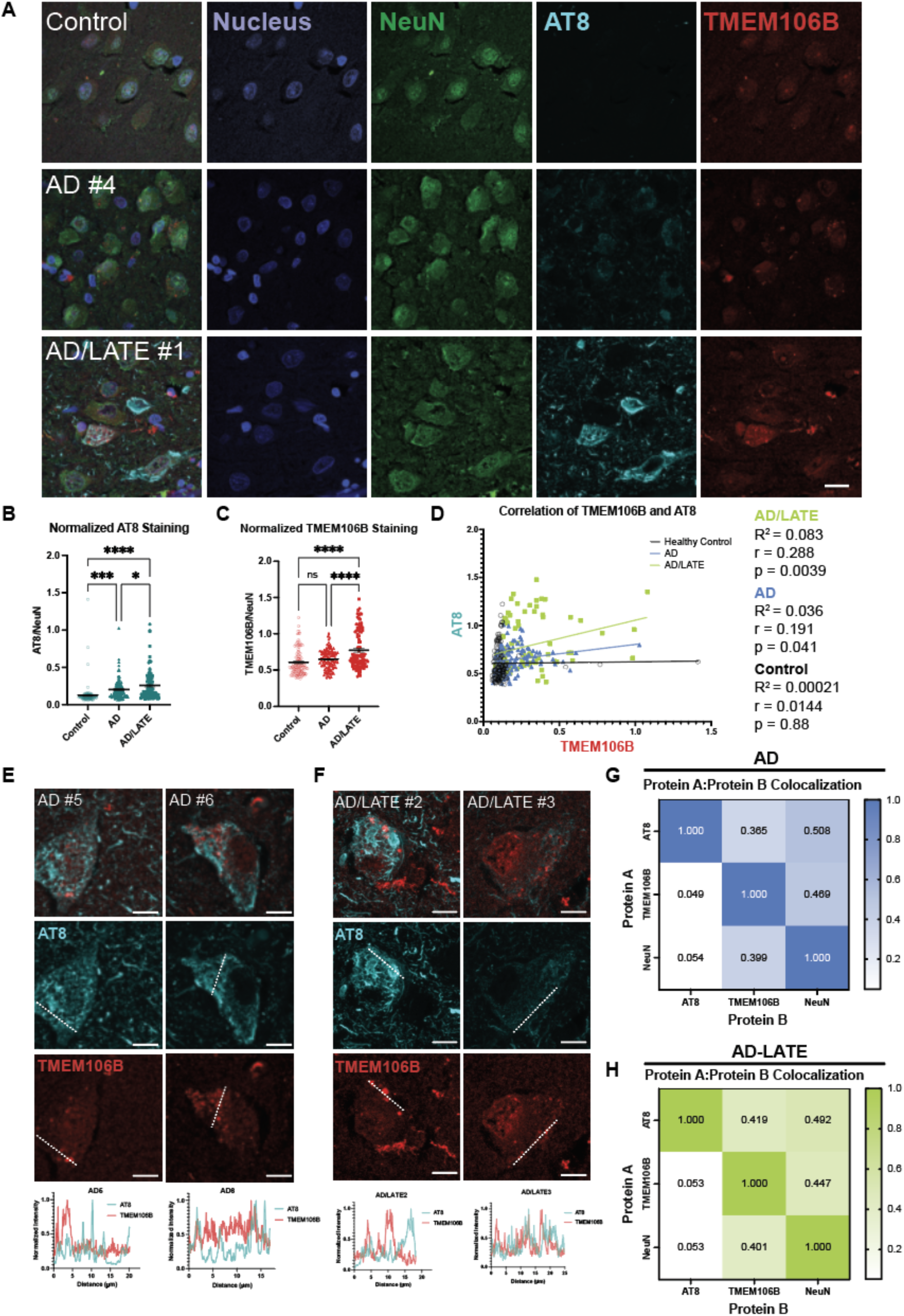
TMEM106B pathology is positively correlated with tau pathology in Alzheimer’s disease (AD) and AD with limbic-predominant age-related TDP-43 encephalopathy (LATE). **(A)** Maximum projection images of human hippocampal tissue stained for DAPI, NeuN, phosphorylated Tau (pTau; AT8), and TMEM106B (Sigma). Scale bar = 20 µm. **(B)** Quantification of AT8 staining normalized to NeuN staining for control, AD, and AD/LATE patients. Each data point represents one cell, bars represent means ± SEM. At least 12 cells were counted per patient with at least 95 cells counted in total for each cohort, n = 5-6. An ordinary one-way ANOVA with Tukey’s multiple comparisons test was used to compare each group. *, p<0.03; ***, p<0.0007; ****, p<0.0001. **(C)** Quantification of TMEM106B staining normalized to NeuN staining for control, AD, and AD/LATE patients. Each data point represents one cell, bars represent means ± SEM. At least 12 cells were counted per patient with at least 95 cells counted in total for each cohort, n = 5-6. An ordinary one-way ANOVA with Tukey’s multiple comparisons test was used to compare each group. ****, p<0.0001. **(D)** A simple linear regression (R^2^) and a Pearson correlation coefficient (r) of the intensity of AT8 and TMEM106B signal intensity for each cell counted. Each dot represents the AT8 and TMEM106B signal from one cell; dots are colored by patient condition. At least 12 cells were counted per patient with at least 95 cells counted in total for each cohort, n = 5-6. Line scans of intensity in patients with AD **(E)**, or AD/LATE **(F)** indicate that the AT8 and TMEM106B signals are not colocalized. The highest signal for each channel was set to 1 and used to normalize all other values. Scans were performed over the white dashed line shown in the images for the individual channels. Heat maps showing the results of rank weighted correlation (RWC) analysis on AD tissues **(G)** and AD-LATE tissues **(H)**. Each cell of the heatmap shows the average RWC coefficient for protein A colocalizing with protein B. A RWC coefficient of 1 indicates perfect colocalization. Four images were analyzed per patient for 5 patients.

We then analyzed the degree of co-localization between AT8 and TMEM106B staining. We found that TMEM106B does not colocalize with AT8 in either AD or AD/LATE patient tissue (**Figure 5E-G**). Within individual cells from multiple patients, line scans do not show intensity trace patterns consistent with colocalization (**Figure 5E, F**). Moreover, taking an unbiased method for analyzing AT8 and TMEM106B signal colocalization, we find that TMEM106B is equally likely to be colocalized with NeuN as with AT8 for both AD and AD/LATE populations (**Figure 5G, H**). Similarly, the reciprocal measure for AT8 reveals that in AD and AD/LATE, AT8 does not colocalize with TMEM106B to a greater extent than it does with NeuN (**Figure 5G, H**).

Taken together, these findings reaffirm prior studies describing TMEM106B aggregation in human tauopathies [64]. Additionally, our data showed that in both PS19 murine model and postmortem tissues from AD and AD/LATE patients, phosphorylated tau burden correlated positively with TMEM106B immunoreactivity, possibly suggesting a conserved role that TMEM106B plays in tauopathy.

## Discussion

TMEM106B, a lysosomal/late endosomal protein originally described as a risk factor for FTD-TDP, has been linked to various neurodegenerative disorders [9, 19, 38, 75, 90]. To date, several *in vivo* models have been used to understand both the function of TMEM106B as well as its potential role as a disease modifier. For example, one recent study showed that TMEM106B knockdown is neuroprotective in both *in vitro* and *in vivo* Parkinson’s disease models, and another study found that loss of TMEM106B exacerbates tau pathology and neurodegeneration in an FTD model [20, 49]. However, these models rely on either overexpression or a knockdown/knockout approach, potentially leading to artificial phenotypes. In this study, we compare the phenotype of endogenous TMEM106B in different disease models and human disease to more accurately understand how TMEM106B pathology relates to neurodegeneration.

As genetic variants in *TMEM106B* have been identified as modifiers of FTLD-TDP in patients with pathological G_4_C_2_ hexanucleotide expansion in *C9orf72* [16, 26, 52, 89–91], we wanted to test whether a C9-mouse model expressing 149 repeats of the disease-associated G_4_C_2_ sequence showed any differences in endogenous TMEM106B localization or expression compared to control animals expressing a 2 repeat control. Intriguingly, we observed novel TMEM106B-positive perinuclear inclusions specifically in AAV-(G_4_C_2_)_149_-injected animals, but not in the control (G_4_C_2_)_2_ mice at 9-months of age (**Figure 1A-E**). These inclusions did not co-localize with markers of autophagy, stress granules, or lysosomes, suggesting these structures do not reflect canonical functioning of TMEM106B or represent bulk degradation of intracellular waste.

One of the hallmark phenotypes of ALS and FTD is the loss of nuclear TDP-43 [2, 7, 51, 79]. Thus, we also examined TDP-43 distribution in the C9 mouse model. We found that cells with TMEM106B inclusions had an aberrantly low TDP-43 N/C ratio (**Figure 1G, H**), suggesting that TMEM106B inclusion formation may be related to TDP-43 mislocalization.

To evaluate whether our findings in a mouse model were relevant to human pathology, we next compared our results in mice to human C9-ALS and C9-ALS/FTD patients. This comparison revealed important differences in pathology. Namely, we did not observe large perinuclear inclusions in any human cells analyzed. Additionally, we did not identify a difference in the percentage of cells with TMEM106B inclusions by diagnosis in humans (**Figure 2C**). However, we did see a disease-dependent correlation between TMEM106B inclusion formation and pathological TDP-43 nuclear clearance (**Figure 2E, F**). Thus, in both the C9-AAV mouse model and C9-ALS human tissue, cells that have TMEM106B positive punctate-like structure have a decreased TDP-43 nuclear to cytoplasmic ratio, suggesting a potential relationship between abnormal TMEM106B pathology and TDP-43 mislocalization in C9-ALS and C9-ALS/FTD in humans.

We next wondered whether the TMEM106B phenotype we observed in C9-animals was present in other models of ALS. As many as 20% of fALS cases are linked to mutations in SOD1 [6], however whether SOD1-related fALS and sALS share common patho-mechanisms is a matter of debate as SOD1-ALS cases with cytoplasmic TDP-43 inclusions are exceptionally rare [86]. Nevertheless, a previous study has reported mislocalization of TDP-43 in end stage SOD1 G93A transgenic mouse model [77]. Furthermore, a recent investigation found age-dependent changes in C-terminal TDP-43 in the spinal cord tissue of SOD1 G93A mouse model as well as in iPSC-derived motor neurons from a SOD1 G17S ALS patient [36].

Here, we analyzed brain and spinal cord tissue from 3-month-old control or transgenic SOD1 G93A mice. Although we observe late-stage pathology, as indicated by the presence of vacuolization, increased SOD1 staining (**Figure 3A, C**) and increased Iba1 staining (**Figure 3B**), we did not observe any changes in TMEM106B staining within the affected brain regions or spinal cord (**Figure 3D, E).** Indeed, in both non-transgenic control and SOD1 G93A-expressing animals, we observed a punctate-like staining of TMEM106B in the midbrain/hindbrain and medulla of the brain, and in the ventral horn region of the spinal cord (**Figure 3D**). Thus, the TMEM106B pathology we observe in C9-ALS does not generalize to all types of ALS and adds to the body of literature that suggests SOD1 ALS is pathophysiologically distinct from sporadic and C9-ALS [14]. However, investigations of endogenous TMEM106B pathology in other populations of ALS and ALS/FTD in which TDP-43 pathology is well-established is an important future area of research.

We next characterized a murine model of tauopathy in which mice transgenically express human tau bearing the dementia-related P301S mutation (PS19) [95]. Consistent with prior reports, we find that 12-month-old mice have significant neuron loss and accumulation of phosphorylated tau (**Figure S4A, B**). However, we did not see significant global changes to TMEM106B levels or localization (**Figure S4C-E**). Moreover, pTau burden was not correlated with TMEM106B intensity (**Figure S4F**). Indeed, AT8-positive aggregates did not colocalize with TMEM106B puncta at the 12-month time point (**Figure S4G**). However, at 9 months of age, when neuronal loss is less severe than at 12 months (**Figure 4A, B**) but significant pTau aggregation has accumulated (**Figure 4C**), PS19 mice have elevated levels of TMEM106B staining, and this increase is positively correlated with pTau burden (**Figure 4D, E).**

To explore whether the colocalization of TMEM106B and pTau generalized to human tissue, we next characterized human AD and AD/LATE tissues. As expected, TMEM106B forms aggregates in these diseases (**Figure 5A, S5A, B**). Moreover, AD/LATE tissues have higher immunoreactivity for TMEM106B (**Figure 5C**), and TMEM106B staining is positively correlated with pTau staining for both AD and AD/LATE (**Figure 5D**). However, in contrast to the PS19 model, TMEM106B does not colocalize with pTau in either AD or AD/LATE (**Figure 5E-H**). Nevertheless, we find a consistent correlation between TMEM106B and pTau burden in both human disease and an *in vivo* model of tauopathy. Our findings warrant additional investigation into whether the correlation between TMEM106B and pathological tau is due to a direct relationship or can be attributed to a common, upstream mechanism.

Given that our findings in both the AAV-induced C9 model and genuine human cases of C9-ALS and ALS/FTD suggest that TMEM106B inclusions are correlated with TDP-43 mislocalization, one important question for future research is whether a similar relationship occurs in other diseases where TDP-43 pathology is present. Indeed, TDP-43 pathology, like TMEM106B aggregation, is not unique to ALS or ALS/FTD; TDP-43 cytoplasmic mislocalization and aggregation has been observed in Alzheimer’s disease and other types of dementia [8, 32, 43, 55], as well as in cognitively normal aged populations [58]. Moreover, previous studies have found that tau burden is related to TDP-43 pathology in AD and other tauopathies [46, 85]. In this work, we found that pTau burden was correlated with TMEM106B intensity at the level of individual cells. Thus, characterizing the relationship between tau, TMEM106B, and TDP-43 in healthy and disease states will be an important step toward describing the nature of these diseases.

It is still unclear whether the TMEM106B aggregates that have been described in neurodegenerative disease and normal aging are a cause or consequence of cellular injury or death. Indeed, the fact that TMEM106B aggregation occurs in healthy aging suggests that, at least to some extent, TMEM106B aggregation is tolerated. On the other hand, the genetic association between TMEM106B and disease, and the identification of disease-associated risk alleles in the *TMEM106B* gene that are associated with increased TMEM106B expression, indicate a relationship between the protein and cell death. In these studies, we characterize endogenous TMEM106B expression and localization in both mouse models of disease and genuine human disease to show distinct phenotypes. Broadly, our results reinforce the importance of relating findings from *in vivo* models to human tissue to optimize translatable research output. Specific to TMEM106B, we find that for C9-related diseases and tauopathies, TMEM106B pathology is correlated with standard measures of disease (i.e., TDP-43 nuclear clearance and pTau accumulation), but that there is no relationship between SOD1 pathology and TMEM106B. Taken together, our findings provide substantial evidence for further investigation into a potential mechanistic link between TMEM106B aggregation and pathological processes in neurodegeneration.

## Methods

### Animals

C57BL/6J were purchased from Jackson Laboratories (Strain #000664) at 4-8 weeks of age. PS19 mice with C57BL/6J background were purchased from Jackson Laboratories (Strain # 024841) at 4-8 weeks of age. SOD1 G93A mice were purchased from Jackson Laboratories (Strain # 002726). At 6-9 weeks of age breeding pairs were established to produce pups for all subsequent experiments. Mice were housed in a constant 14-hour light/10-hour dark cycle and allowed access to food and water *ad libitum*. In this study, tissue from 8 (G_4_C_2_)_2_ repeat and 9 (G_4_C_2_)_149_ repeat injected animals, 7 SOD1 G93A (6 females, 1 male) along with 8 non-transgenic controls (4 females, 4 males), 5 9-month-old PS19 along with 5 non-transgenic controls, and 5 12-month-old PS19 along with 3 non-transgenic controls were used. All animal procedures complied with animal protocols approved by the Animal Use Committee at the Johns Hopkins University School of Medicine (JHUSOM).

### Neonatal Viral Injections

The AAV2/9-(G_4_C_2_)_2_ and AAV2/9-(G_4_C_2_)_149_ viruses were provided by Dr. Leonard Petrucelli at Mayo Clinic Jacksonville. The viruses were prepared as previously described [12, 82]. AAV viral aliquots were thawed on ice and spun down in a centrifuge at 4°C. In a sterile hood, viruses were diluted to 1.5×10^10^ viral genomes/µL (vg/µL) with sterile PBS and were stored on ice until time of injection. Intracerebroventricular (ICV) injections of AAV were performed on C57BL/6J postnatal day 0 (P0) pups. AAV dilutions were prepared on the day of injections. Pups underwent cryoanesthesia on ice for approximately 3 minutes or until pups exhibited no movement. A 32-gauge needle (Hamilton; Small RN 32 gauge, 0.5 inch needle, point style 4) attached to a 10 µL syringe (Hamilton, Model 701 RN) was inserted approximately two fifths of the distance between the lambda and each eye at a 30° angle from the surface of the head and was held at a depth of about 2 mm. 2 µL of virus was manually injected into each cerebral ventricle and the needle was held in place for an additional 5 seconds after each injection to prevent back flow. After injections, pups were placed on a heating pad until fully recovered and then returned to their home cages with the dam. Any pups with back flow from the injection were excluded from the study.

### Tissue Harvesting

#### Brain

The anesthetized mouse was transcardially perfused with ice-cold PBS containing 10 U/mL heparin (Sigma-Aldrich H3149) for approximately 5 minutes. Subsequently, the brain was removed and put in a conical tube containing 4% paraformaldehyde in PBS overnight at 4°C and then moved into PBS containing 0.1% sodium azide for long store storage.

#### Spinal Cord

The spinal cord was dissected, similar to previously described [39]. Laminectomy was performed by gentle cutting laminae at 3 and 9 o’clock directions from cervical to lumbar level. The lumbar vertebra was cut out. The spinal nerve roots were gently cut off on both sides and the spinal cord was removed from the spinal canal. The spinal cord was then cut into Thoracic and Lumbar sections. Both Thoracic and Lumbar sections were put into microcentrifuge tubes containing 4% paraformaldehyde in PBS overnight at 4°C and then moved into PBS containing 0.1% sodium azide for long term storage.

### Immunofluorescence

All mouse brain tissue were paraffin embedded and cut in sagittal orientation with 5 µm thickness. Mouse spinal cord tissue was also paraffin embedded and cut in cross section orientation with 5 µm thickness. Formalin-fixed-paraffin-embedded (FFPE) sections were deparaffinized in xylene and rehydrated through a series of ethanol solutions. Antigen retrieval was performed in 10 mM sodium citrate buffer, pH 6.0 for 60 minutes in a steamer and then allowed to cool for 10 minutes. Following washing with deionized water and PBS, the tissue was permeabilized with 0.2% (mouse tissue) or 0.4% (human tissue) Triton X-100 in PBS for 10 minutes at room temperature. The sections were then washed with PBS with 0.05% Tween (PBST; mouse) or PBS (human) 3x. Mouse tissues were blocked with 10% Normal Goat serum containing 0.05% Tween for 1 hour at room temperature; human tissues were blocked in DAKO protein-free serum block (DAKO X0909) overnight at 4 °C.

Mouse sections were immunostained with primary antibodies (**Table 1**) diluted in blocking buffer overnight at 4 °C and were subsequently washed with PBST (PBS with 0.05% Tween). Secondary antibodies diluted in blocking solution were incubated at room temperature for 1 hour. After secondary antibody staining, the sections were processed with the autofluorescence eliminator reagent (Millipore Sigma #2160) according to the manufacturer’s instructions.

Sections were then incubated with PBST and Hoechst (1:1000) for 10 minutes, followed by additional washes with PBST. Slides were mounted on a coverslip with ProLong Gold mounting solution (ThermoFisher Scientific P36931).

Human tissues were immunostained with primary antibody diluted in DAKO antibody diluent (DAKO S0809) and stored in a humidified chamber at 4 °C for 2-3 days. Before applying secondary antibody, tissues were brought to room temperature for ∼20 minutes, then washed 3x with PBS. Secondary antibodies were diluted in DAKO antibody diluent and applied to slides for 1 hour at room temperature. After secondary antibody staining, the sections were processed with the autofluorescence eliminator reagent (Millipore Sigma #2160) according to the manufacturer’s instructions. Tissues were then washed 3x with PBS, stained with DAPI, and washed another 2x with PBS before mounting with ProLong Gold.

### DAB Staining

FFPE sections were deparaffinized in xylene and rehydrated through a series of ethanol solutions. Antigen retrieval was performed in 10 mM sodium citrate buffer, pH 6.0 for 1 hour. Tissues were immunostained with primary antibodies overnight. DAKO Envision+HRP polymer kits (K4003 and K4001) were used, and the reaction was visualized using ImmPACT® VIP Substrate Kit (Vector Laboratories SK-4605). Sections were then counterstained with Hematoxylin QS Counterstain (Vector Laboratories H-3404-100) and mounted with Aqua-Poly/Mount (Polysciences 18606-20).

### Microscopy

For DAB, slides were imaged as 20x magnification tiles or individual 63x magnification images using Zeiss Axio Imager. For IF, slides were imaged at 20x, 40x, or 63x magnification using a Zeiss LSM 980 with Airyscan, as indicated.

### Human postmortem tissue

All human ALS tissues used within this study were obtained from Dr. Alyssa Coyne (Johns Hopkins), Dr. Dennis Dickson (Mayo Clinic), and the Target ALS Postmortem Tissue Core (see Table 3). All human AD tissues used within this study were obtained from the Johns Hopkins Alzheimer’s Disease Research Center (IRB 00082277, see Table 4).

**Table 3.** Human tissue demographics – C9ORF72.

| Patient ID | Condition | Age | Sex |
| --- | --- | --- | --- |
| 001 (1) | Control | 52 | M |
| 005 | Control | 72 | M |
| 008 | Control | 71 | F |
| 002 (1) | C9-ALS | 61 | F |
| 003 (2) | C9-ALS | 54 | F |
| 004 | C9-ALS/FTD | 74 | M |
| 006 | C9-ALS | 56 | F |
| 007 | C9-ALS/FTD | 68 | F |
| 009 | C9-ALS | 47 | M |
| 010 | C9-ALS | 62 | M |

*Numbers in parentheses refer to the patient numbering convention used in figures.*

**Table 4.**
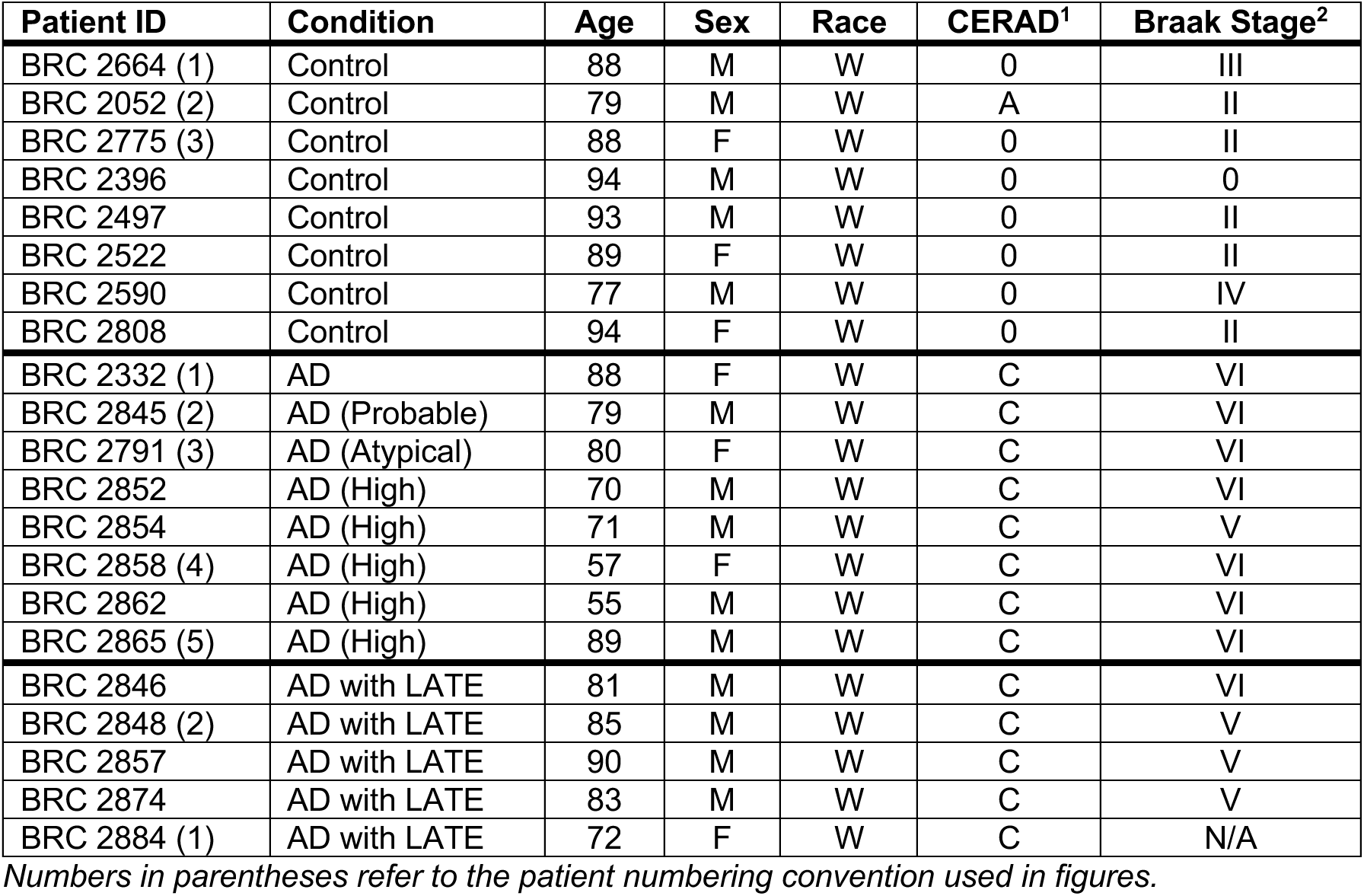
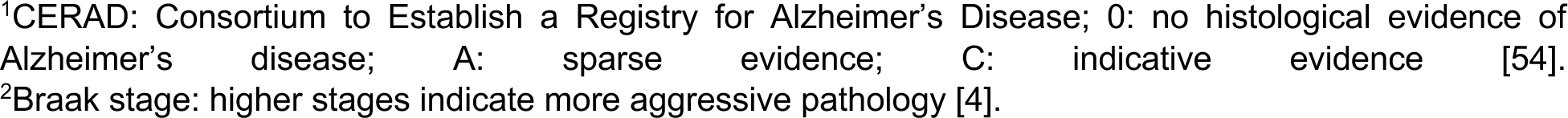
Human tissue demographics – Alzheimer’s Disease.

| Patient ID | Condition | Age | Sex | Race | CERAD<sup>1</sup> | Braak Stage<sup>2</sup> |
| --- | --- | --- | --- | --- | --- | --- |
| BRC 2664 (1) | Control | 88 | M | W | 0 | III |
| BRC 2052 (2) | Control | 79 | M | W | A | II |
| BRC 2775 (3) | Control | 88 | F | W | 0 | II |
| BRC 2396 | Control | 94 | M | W | 0 | 0 |
| BRC 2497 | Control | 93 | M | W | 0 | II |
| BRC 2522 | Control | 89 | F | W | 0 | II |
| BRC 2590 | Control | 77 | M | W | 0 | IV |
| BRC 2808 | Control | 94 | F | W | 0 | II |
| BRC 2332 (1) | AD | 88 | F | W | C | VI |
| BRC 2845 (2) | AD (Probable) | 79 | M | W | C | VI |
| BRC 2791 (3) | AD (Atypical) | 80 | F | W | C | VI |
| BRC 2852 | AD (High) | 70 | M | W | C | VI |
| BRC 2854 | AD (High) | 71 | M | W | C | V |
| BRC 2858 (4) | AD (High) | 57 | F | W | C | VI |
| BRC 2862 | AD (High) | 55 | M | W | C | VI |
| BRC 2865 (5) | AD (High) | 89 | M | W | C | VI |
| BRC 2846 | AD with LATE | 81 | M | W | C | VI |
| BRC 2848 (2) | AD with LATE | 85 | M | W | C | V |
| BRC 2857 | AD with LATE | 90 | M | W | C | V |
| BRC 2874 | AD with LATE | 83 | M | W | C | V |
| BRC 2884 (1) | AD with LATE | 72 | F | W | C | N/A |

*Numbers in parentheses refer to the patient numbering convention used in figures.*
<sup>1</sup>CERAD: Consortium to Establish a Registry for Alzheimer's Disease; 0: no histological evidence of Alzheimer's disease; A: sparse evidence; C: indicative evidence [54]. <sup>2</sup>Braak stage: higher stages indicate more aggressive pathology [4].

### C9 Tissue Quantification

The motor cortex was imaged at 63x using confocal microscopy. Maximum intensity projections were used for all quantification purposes. The number of TMEM106B inclusions and total number of neurons in the field were manually counted. TDP-43 N/C ratio was quantified in ImageJ by manually drawing a region with the guide of Hoechst channel as a nuclear mask and NeuN channel as a cytoplasm mask.

### Mouse PS19 Tissue Image Quantification

For TMEM106B and AT8 staining intensity in the PS19 studies, 20x images were taken of the dentate gyrus. A mask was drawn using ImageJ around the NeuN+ cells in these regions. The mean intensity of NeuN, TMEM106B, and AT8 were then quantified in ImageJ.

The signal of TMEM106B and AT8 were each normalized to the respective NeuN signal for each image. For each image, the NeuN-normalized signal for TMEM106B and AT8 of each image was used for correlation analysis. Colocalization analysis was measured across the entire image for maximum intensity projections of the NeuN, AT8, and TMEM106B channels using MeasureColocalization in CellProfiler with the threshold as percentage of maximum intensity set to 15.0. The Rank Weighted Colocalization (RWC) coefficient was used, as this measure incorporates image intensity by first ranking each pixel in each image from 1 – n based on intensity, with 1 representing the highest intensity pixel [78]. The RWC coefficient is calculated using the following equations:

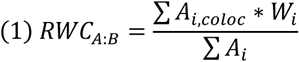

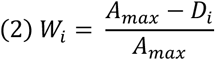

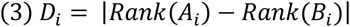

Where *A_i_* is the intensity of image A at a given pixel, and A_i, coloc_ = A_i_ if B_i_ is > 0, and A_i, coloc_ = 0 if B_i_ is < 0 (i.e., A_i, coloc_ represents only pixels where images A and B both have a positive signal). W_i_ is weight, which incorporates A_max_ as the maximum rank of the pixel in either A and B, and D_i_, the absolute value of the difference between the rank of the pixel for each image.

### Human AD Tissue Image Quantification

Cells in the CA1 region of the hippocampus were imaged at 63x using confocal microscopy. Maximum intensity projections were used for all quantification purposes. To quantify target intensity, a cellular mask was manually drawn using NeuN; this mask was then used to quantify the mean intensity of NeuN, TMEM106B, and AT8. TMEM106B and AT8 intensities were normalized to the NeuN intensity for each cell. For line scans, a line was drawn over cells using ImageJ and a plot profile was generated for both the TMEM106B and AT8 channels. These values, paired by position along the line, were then plotted for each cell.

## Contributions

A.D., S.C.A., C.M.F., and L.R. designed the experiments. A.D., S.C.A., C.M.F., S.V., and L. M., conducted all in-life experimentation. A.D., S.C.A., C.M.F., and J.L. performed all molecular and histological experimentation. A.D., S.C.A., C.M.F., and J.D.R. performed all data analysis. A.D., S.C.A., C.M.F. wrote the paper. All authors reviewed and edited this manuscript.

## Acknowledgments

We thank all the patients and their families who have donated their tissue for use in science. This work was funded by NIH-NINDS (R01 5R35NS132179-02, J.D.R.) and the ALS Association Milton Safenowitz Postdoctoral Fellowship (S.C.A.). We would like to especially thank the Johns Hopkins Alzheimer’s Disease Research Center (National Institutes on Aging grant P50AG005146) for providing human AD tissue, Dr. Alyssa Coyne, Dr. Dennis Dickson, and the Target ALS Postmortem Tissue Core for providing human ALS tissue used in this study, and Johns Hopkins University undergraduate student Katie Koo for assistance in this project.

## Supplemental Figures

**Figure S1:**
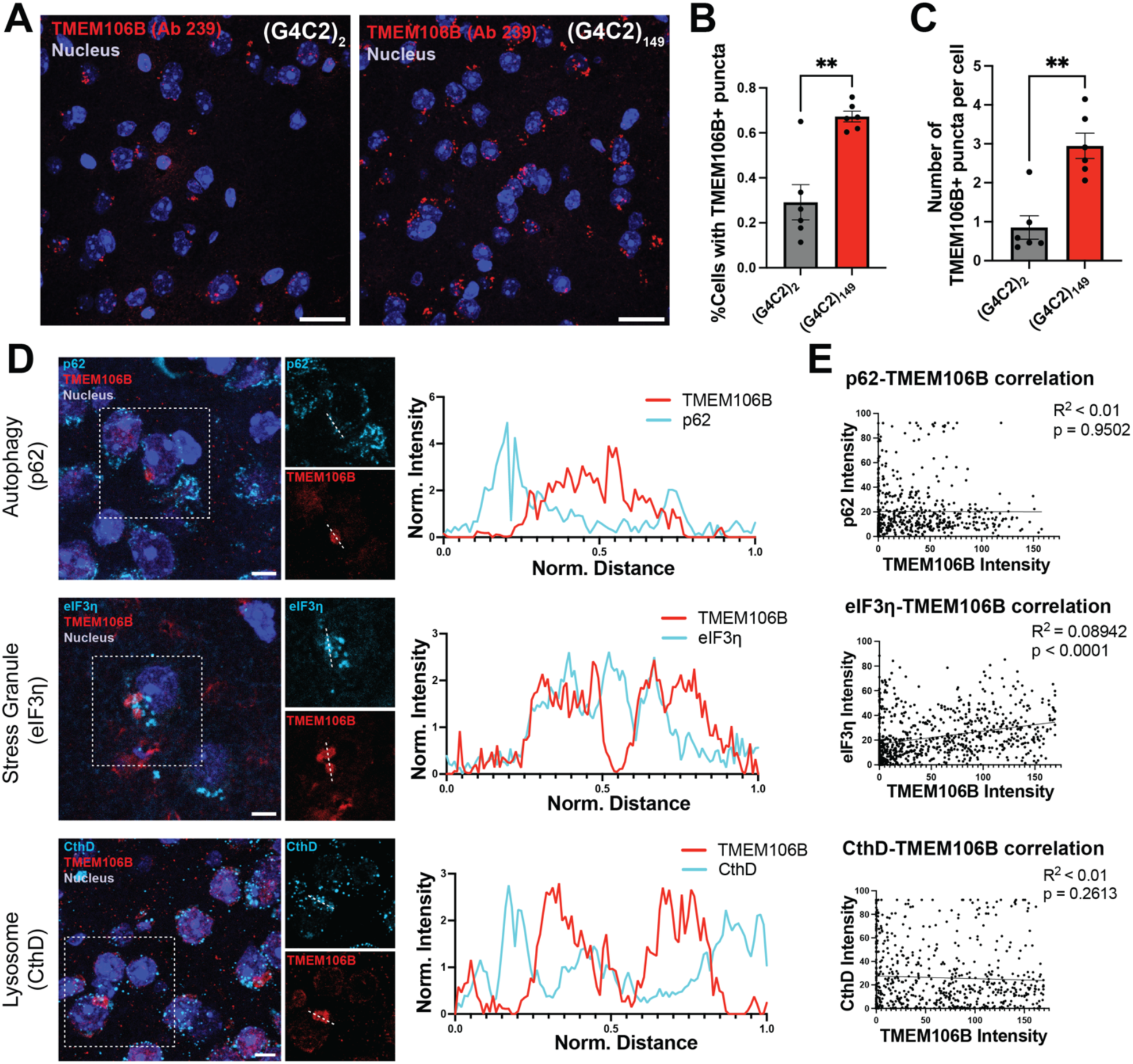
TMEM106B immunoreactivity probed with the TMEM239 antibody and co-localization analysis of TMEM106B inclusion with other cellular markers. **(A)** Representative immunofluorescence images of the motor cortices of (G_4_C_2_)_2_ (n = 6) and (G_4_C_2_)_149_ (n = 6) mice probed by the TMEM239 antibody. Scale bar = 20 µm. **(B)** Quantification of the percentage of cells with TMEM106B puncta probed by the TMEM239 antibody. Dots represent individual animals, bars represent means ± SEM. Unpaired Welch’s t test, p=0.0036. **(C)** Quantification of the average number of TMEM106B puncta probed by the TMEM239 antibody. Mann-Whitney test, p=0.0043. **(D)** Representative images of TMEM106B perinuclear inclusion probed with the TMEM-Sigma antibody co-stained with an autophagy marker (p62), a stress granule marker (eukaryotic initiation factor 3η, eIF3η), and a lysosome marker (cathepsin D, CthD). A line crossing through the TMEM106B inclusion is shown on the left panel and the normalized intensity of TMEM106B and each marker along the line are plotted on the right panel. **(E)** Correlation analysis reveals that TMEM106B inclusions are not associated with the lysosome or autophagic bodies, but are positively correlated with the presence of stress granules. Each data point stands for a pixel on a line crossing through TMEM106B inclusions.

**Figure S2:**
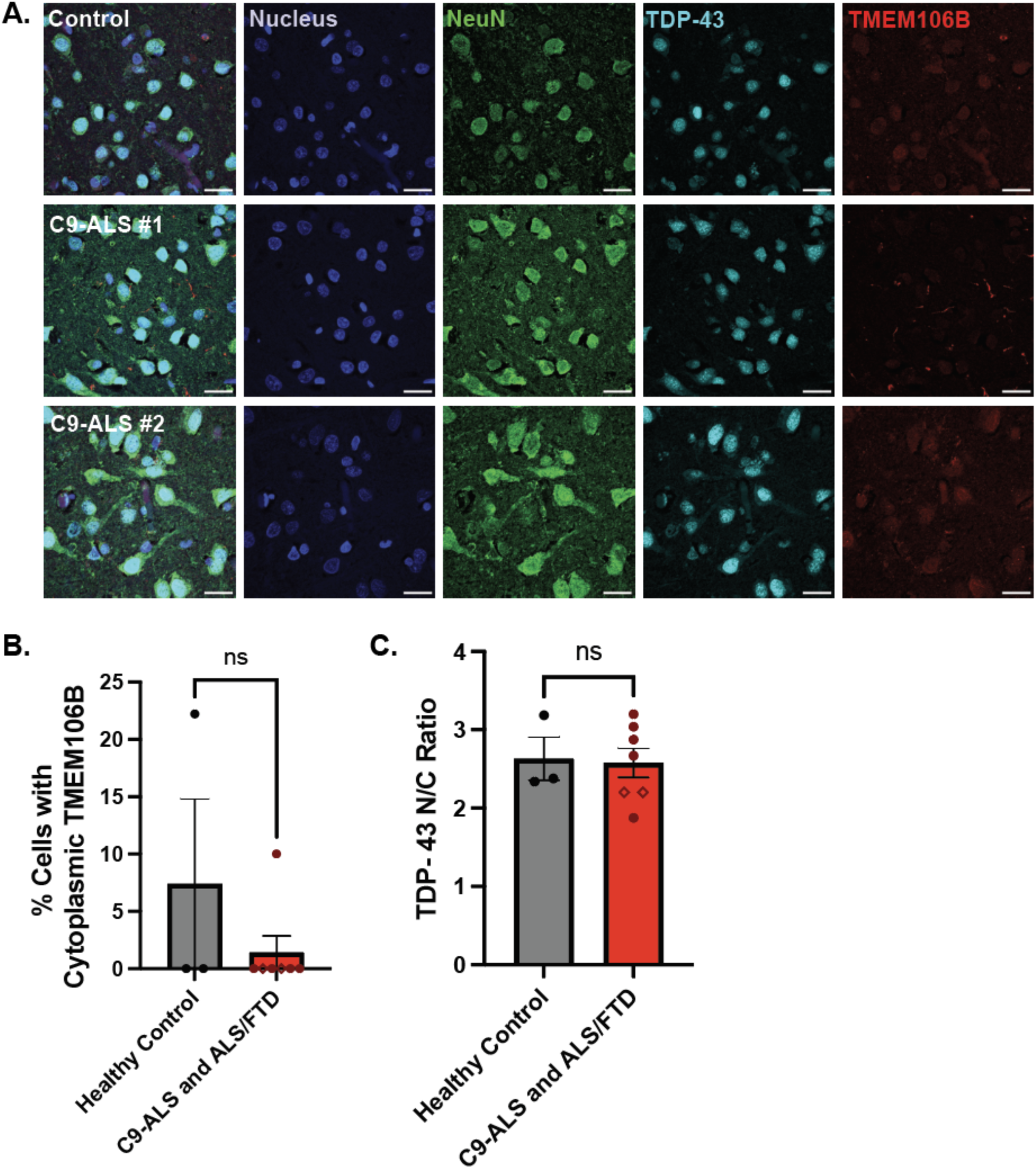
TMEM106B pathology is not observed in the occipital lobe of C9-ALS or ALS/FTD patients. **(A)** Representative immunofluorescence images of human occipital cortex co-stained for NeuN, TDP-43, and TMEM106B (Sigma antibody). The patients shown here are the same as shown in Figure 2. Scale bar = 20 µm. **(B)** Quantification of the percent of cells with cytoplasmic TMEM106B puncta shows there is no difference between control and disease. For the C9-ALS and ALS/FTD bars, closed dots represent C9-ALS, and open diamonds represent C9-ALS/FTD. Data points represent averages from individual people (healthy control n = 3, C9-ALS and ALS/FTD n = 7), bars represent means ± SEM. A t-test was used to compare groups. **(C)** Quantification of the TDP-43 nuclear to cytoplasmic (N/C) ratio in neurons shows there is no difference between control and disease. For the C9-ALS and ALS/FTD bars, closed dots represent C9-ALS, and open diamonds represent C9-ALS/FTD. Data points represent averages from individual people (healthy control n = 3, C9-ALS and ALS/FTD n = 7), bars represent means ± SEM. A t-test was used to compare groups.

**Figure S3:**
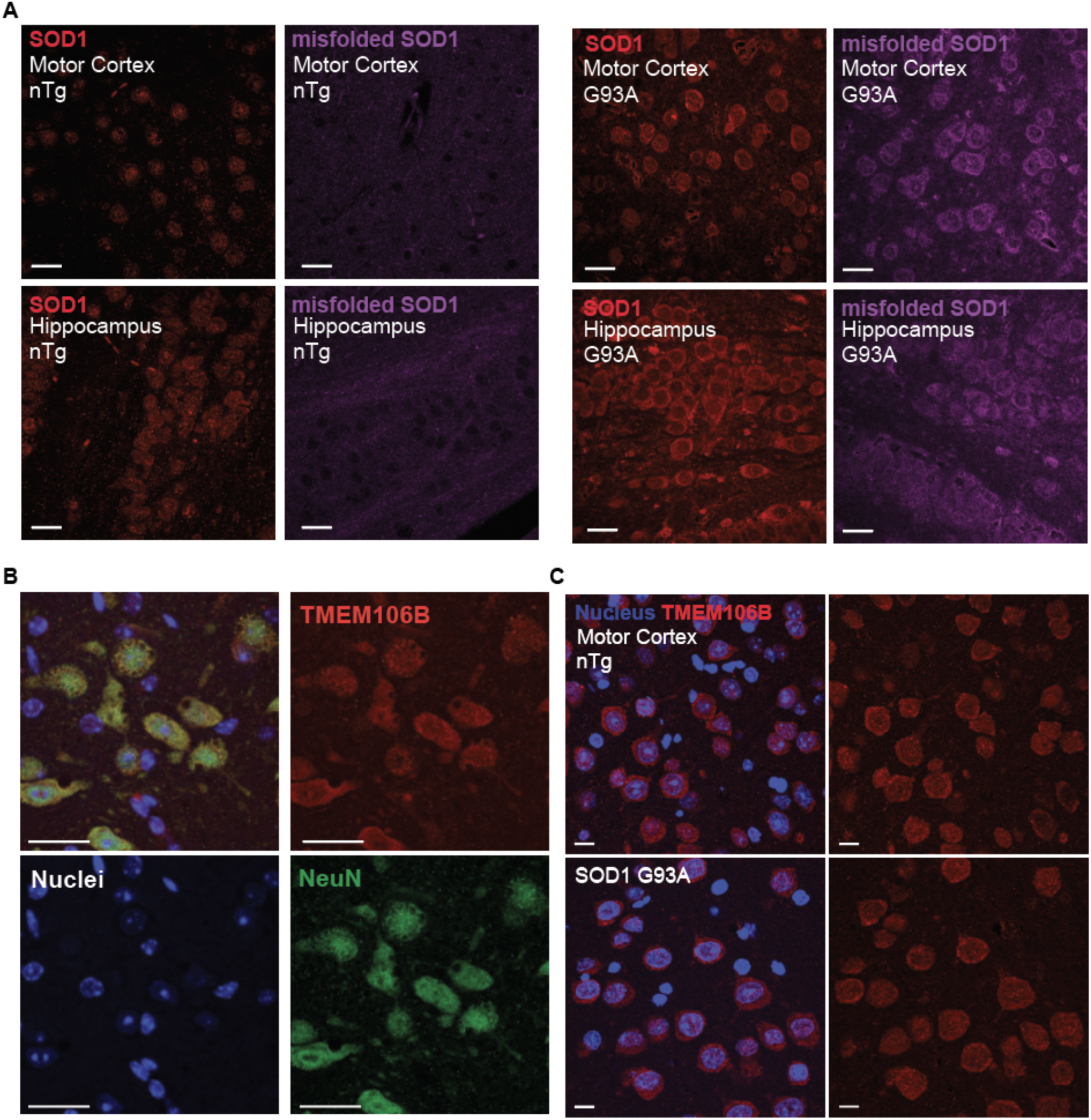
TMEM106B localization in the motor cortex is consistent between control and transgenic SOD1 mice. **(A)** In addition to the midbrain and hindbrain, we observe SOD1 overexpression in the motor cortex and hippocampus of SOD1 transgenic mice. We also observe positive staining for an antibody that recognizes misfolded SOD1. Images are representative from n = 8 non-transgenic (nTg) and 7 transgenic animals expressing the G93A SOD1 mutant protein. Scale bar = 25 µm. **(B)** High-resolution Airyscan microscopy taken of cells exhibiting vacuolization does not show strong TMEM106B signal near or around vacuoles. Scale bar = 25 µm. **(C)** As was observed in the midbrain and hindbrain, TMEM106B staining is indistinguishable in the motor cortex of transgenic and nTg animals. Images are representative from n = 8 nTg and 7 transgenic animals. Scale bar = 10 µm.

**Figure S4:**
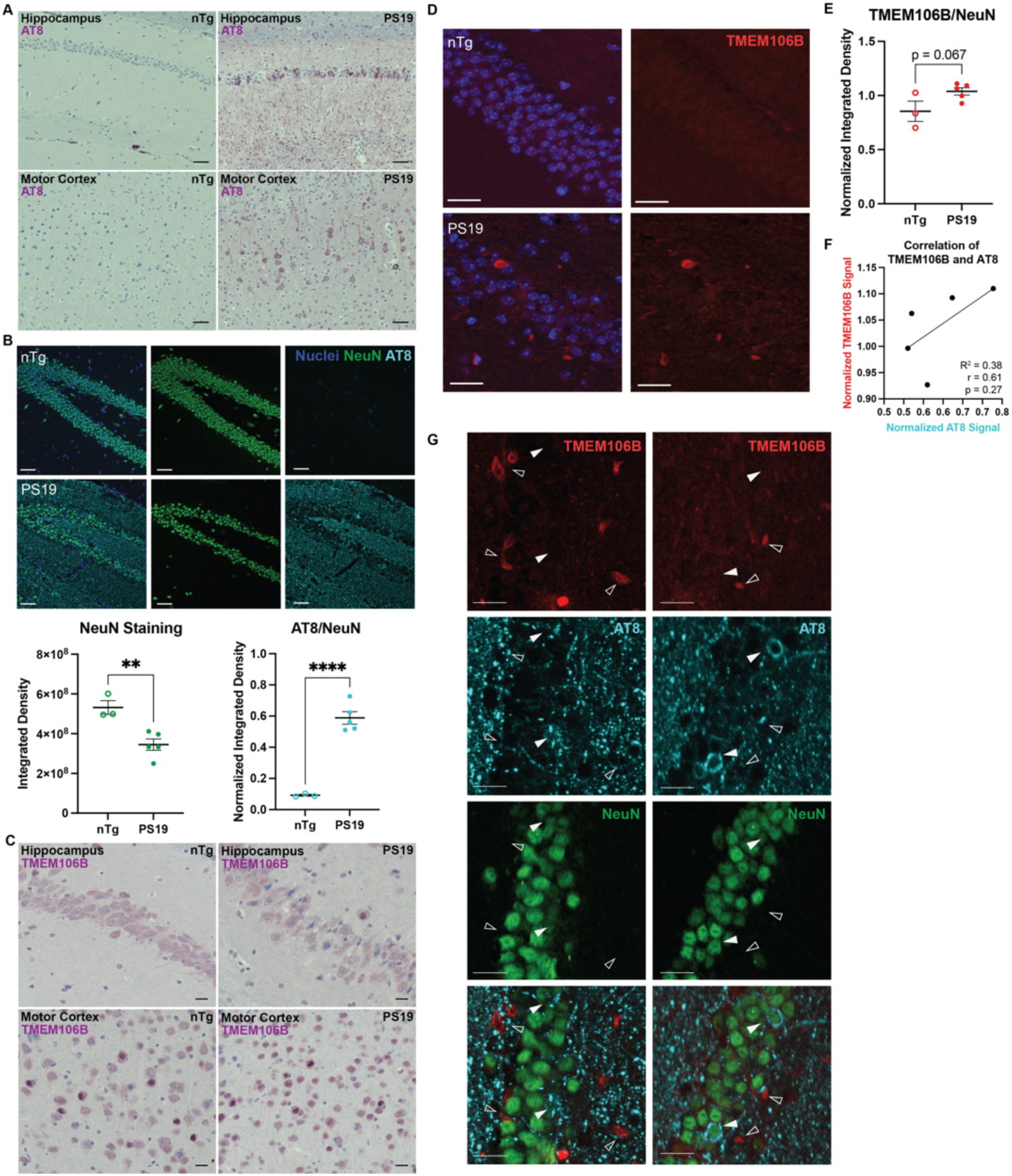
TMEM106B pathology is distinct from phosphorylated tau in PS19 mice in aged animals. **(A)** Representative images of CA1 and motor cortex with DAB staining against phosphorylate tau (AT8) in non-transgenic (nTg) animals and animals expressing the P301S mutant tau (PS19). Scale bar = 50 µm. **(B)** Immunofluorescence confocal microscopy of NeuN and AT8 in the dentate gyrus region of the hippocampus. Scale bar = 50 µm. Below, left: the integrated density signal for NeuN for each image. Below, right: the signal for each AT8 image was normalized to the signal of its corresponding NeuN channel. Each data point represents an individual image from n = 3 nTg animals and 5 PS19 animals. Bars represent means ± SEM. A student’s t-test was used to compare nTg and transgenic animals; **, p<0.007; ****, p<0.0001. **(C)** CA1 and motor cortex with DAB staining against TMEM106B. Scale bar = 20 µm. **(D)** Immunofluorescence confocal microscopy of dentate gyrus hippocampal tissue for non-transgenic and PS19 animals. Scale bar = 25 µm. **(E)** Quantification of TMEM106B intensity normalized to NeuN intensity. Each data point represents an individual image from n = 3 nTg animals and 5 PS19 animals. Bars represent means ± SEM. A student’s t-test was used to compare nTg and PS19 animals. **(F)** Correlation analysis between the TMEM106B signal and AT8 staining; Pearson r value of 0.61, p = 0.27. Linear regression of the data yields a line with an R^2^ value of 0.38 with a slope that does not significantly deviate from zero. Data points represent individual images from n = 5 transgenic animals. **(G)** Images from the dentate gyrus two different PS19 animals showing instances of pTau aggregation (solid arrows) or TMEM106B-positive staining outside of the neuronal cell layer (empty arrows). Scale bar = 20 µm.

**Figure S5:**
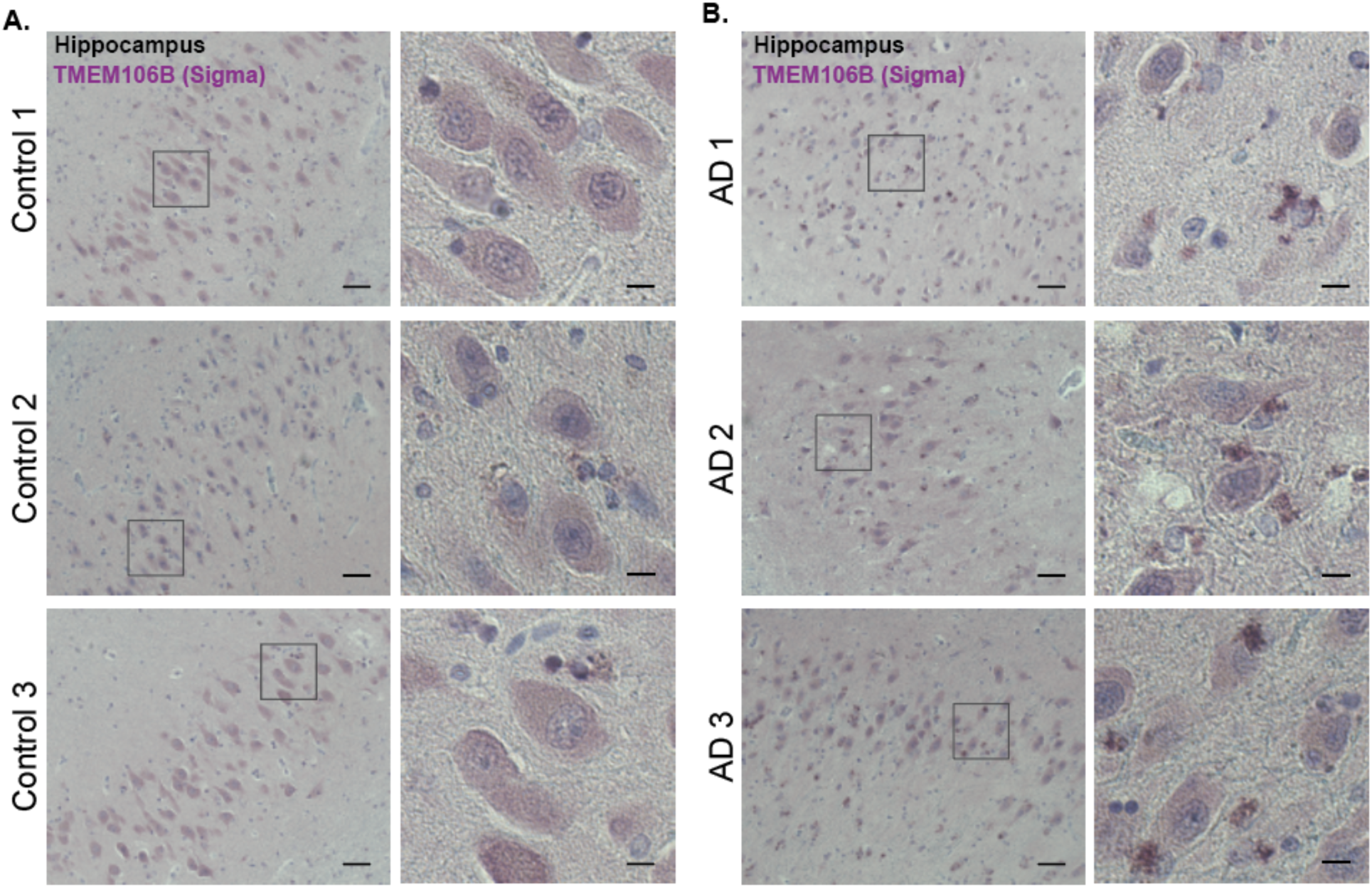
TMEM106B forms extracellular inclusions in human Alzheimer’s disease. **(A)** Representative images of the pyramidal layer with DAB staining against TMEM106B in three neurologically healthy controls. Scale bar = 50 µm, Inset scale bars = 10 µm. **(B)** Representative images of the pyramidal layer with DAB staining against TMEM106B in three AD patients. Scale bar = 50 µm, Inset scale bars = 10 µm.

